# Distal Radius Microstructure and Finite Element Bone Strain Are Related to Site-Specific Mechanical Loading and Areal Bone Mineral Density in Premenopausal Women

**DOI:** 10.1101/146258

**Authors:** Megan E. Mancuso, Joshua E. Johnson, Sabahat S. Ahmed, Tiffiny A. Butler, Karen L. Troy

**Author notes:** Corresponding Author: Karen L. Troy, Ph.D, Department of Biomedical Engineering, Worcester Polytechnic Institute, 100 Institute Road, Worcester, MA 01701.

## Abstract

While weight-bearing and resistive exercise modestly increases aBMD, the precise relationship between physical activity and bone microstructure, and strain in humans is not known. Previously, we established a voluntary upper-extremity loading model that assigns a person’s target force based on their subject-specific, continuum FE-estimated radius bone strain. Here, our purpose was to quantify the inter-individual variability in radius microstructure and FE-estimated strain explained by site-specific mechanical loading history, and to determine whether variability in strain is captured by aBMD, a clinically relevant measure of bone density and fracture risk. Seventy-two women aged 21-40 were included in this cross-sectional analysis. High resolution peripheral quantitative computed tomography (HRpQCT) was used to measure macro- and micro-structure in the distal radius. Mean energy equivalent strain in the distal radius was calculated from continuum finite element models generated from clinical resolution CT images of the forearm. Areal BMD was used in a nonlinear regression model to predict FE strain. Hierarchical linear regression models were used to assess the predictive capability of intrinsic (age, height) and modifiable (body mass, grip strength, physical activity) predictors. Fifty-one percent of the variability in FE bone strain was explained by its relationship with aBMD, with higher density predicting lower strains. Age and height explained up to 31.6% of the variance in microstructural parameters. Body mass explained 9.1% and 10.0% of the variance in aBMD and bone strain, respectively, with higher body mass indicative of greater density. Overall, results suggest that meaningful differences in bone structure and strain can be predicted by subject characteristics.

**Highlights:**

- Areal bone mineral density (aBMD) explains 51% of the variability in bone strain.
- Adult bone loading predicts greater cortical porosity and trabecular density.
- Greater body mass predicts greater aBMD and lower bone strain.

## 1. Introduction

Bone is a mechanosensitive tissue, with a complex structure adapted to habitual mechanical loads. Controlled human trials have shown that high-impact [1,2] and resistive [3,4] exercises lead to modest but statistically significant increases in areal bone mineral density (aBMD). As a result, The National Osteoporosis Foundation recommends that women perform weight-bearing and muscle-strengthening exercises throughout their lifespan to reduce the risk of osteoporotic fracture [5]. Despite this knowledge, it remains unclear which exercises are most effective at increasing bone strength [5,6], and there are no systematic methods to prescribe loading to specific individuals or clinical populations [7]. This is largely because the precise relationship between bone structure and loading during physical activity in humans remains unknown, constraining the translation of animal work to the clinic.

Mechanical stimuli related to bone strain are understood to drive adaptation of bone structure to loading. In animal models, strain magnitude [8], rate [9], number of loading cycles [10,11], and strain energy density [12,13] modulate bone adaptation. However, the relationship between bone strain and adaptation has not been studied directly in humans due to challenges associated with measuring local tissue loading. Strain can only be measured invasively, using strain gauges applied to the periosteal surface directly or on a bone staple [14,15]. This is limited in that strain is measured only on a small area of the external bone surface for short time periods. Computed-tomography (CT)-based finite element (FE) models have enabled the non-invasive, subject specific estimation of strain throughout a bone volume. Using this technology, we previously established a tunable upper-extremity axial loading model in humans [16], which uses FE-estimated bone strain [17] as a basis for prescribing target forces. The radius was selected because it is the most common site of osteoporotic fractures [18], prescribed loads are not confounded by weight bearing, and the forearm can be imaged at high resolutions [19]. The loading task is simulated using clinical CT-based FE models [17], and the resulting force-strain relationship is used to assign a subject-specific force that generates a desired average radius strain. We have shown that forearm loading magnitude can be voluntarily manipulated to achieve specific strains during this task [16], highlighting its potential to answer several important questions about human bone adaptation.

Radius bone strength and the strain it experiences due to a given force, is highly variable [14] between individuals, even those with similar bone mineral content [17]. Understanding the sources of this variability is an important step towards individualized exercise prescription. Habitual external loads from sports participation are one source of variability. Studies using areal bone mineral density (aBMD) as an outcome have shown that high volumes of physical activity improve bone density, as seen in the forearms of competitive tennis players [20], racquetball players [21], and gymnasts [22,23]. Muscle forces also contribute to the habitual loads experienced by the forearm. Grip strength is related to radius structural indices calculated from peripheral quantitative computed tomography [24,25], but not to ultradistal radius aBMD [26]. Body mass is another source of external loading. Although body mass has been consistently linked to aBMD at weight bearing sites [27–30], conflicting results have been found for the radius [26,31]. Ultimately, it is not known whether loading history is related to radius bone strain in the same way that it is related to aBMD.

Bone microstructure, which can be measured using high-resolution peripheral quantitative computed tomography (HRpQCT) [19,32], is also an important determinant of bone mechanics [33] and fracture risk [34–36]. However, limited data exist regarding the extent to which bone microstructure is modulated by mechanical loading history. One group found that HRpQCT parameters in the radius of young adult males were associated with present and past physical activity [37] and participation in soccer [38], while another study found similar sport-specific differences in soccer playing males but not females [39]. The extent to which loading history affects radius microstructure in females with an average level of upper-extremity loading (i.e. non-athletes) has not been examined.

Here, our primary purpose was to quantify the inter-individual variability in radius microstructure and FE-estimated strain explained by site-specific mechanical loading history. We hypothesized that greater site-specific loading, indicated by high levels of physical activity, grip strength, and body mass would predict favorable bone structure and lower FE strain, independent of age and height. Our secondary purpose was to determine whether variability in strain is captured by aBMD, a clinically relevant measure of bone density and fracture risk.

## 2. Materials and Methods

### 2.1 Participants

Healthy females age 21-40 were recruited from the greater Worcester area as part of a larger, institutionally approved longitudinal experiment (Figure 1). The present study reports baseline cross-sectional data from the parent study. Women responding to online advertisements were contacted and screened via telephone survey. Individuals with irregular menstrual cycles, body mass indices outside the range 18-25 kg/m^2^, no regular calcium intake, or those taking medications known to affect bone metabolism were excluded. Because subjects were being screened for a prospective loading intervention study, individuals with a history of radius fracture or injury of the non-dominant shoulder or elbow, and those regularly participating (> 2 time per month) in sports that apply high-impact loads to the forearm (e.g. gymnastics, volleyball) were also excluded. Those satisfying the initial inclusion criteria were screened for 25-hydroxyvitamin D serum levels and forearm DXA T-score during a prescreening visit (Hologic; Marlborough, MA). DXA scans were performed using the Hologic Discovery C to image the non-dominant forearm according to the manufacturer’s standard protocol, and used to calculate T-score and aBMD within the ultradistal and total forearm regions (Figure 2a). T-scores were used to determine study eligibility, while ultradistal aBMD for enrolled subjects was used in analysis to quantify the relationship between aBMD and FE-estimated strain. Qualified subjects had 25-hydroxyvitamin D serum above 20 ng/ml and a total forearm DXA T-score between −2.5 and 1.0. Data for qualified subjects (n=82) were collected either during the screening or a single visit within approximately two weeks of screening. All participants provided written, informed consent between January 2014 and November 2016.

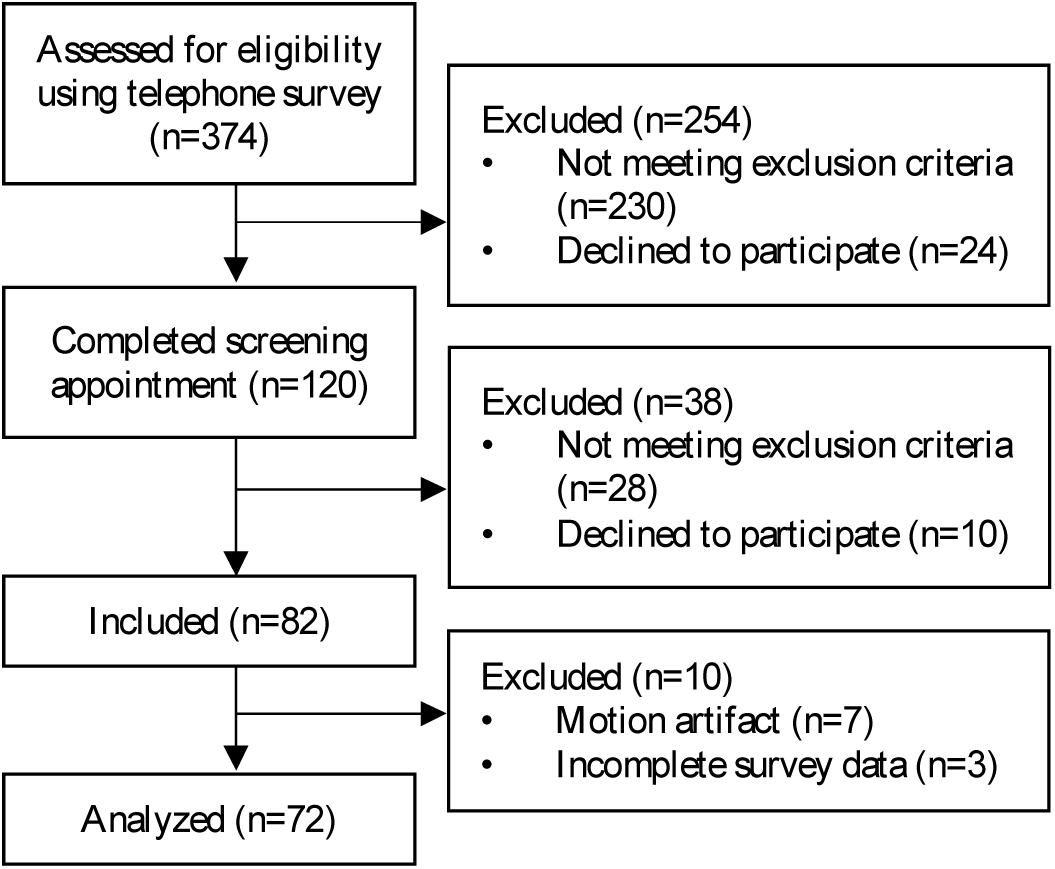

**Figure 2:**
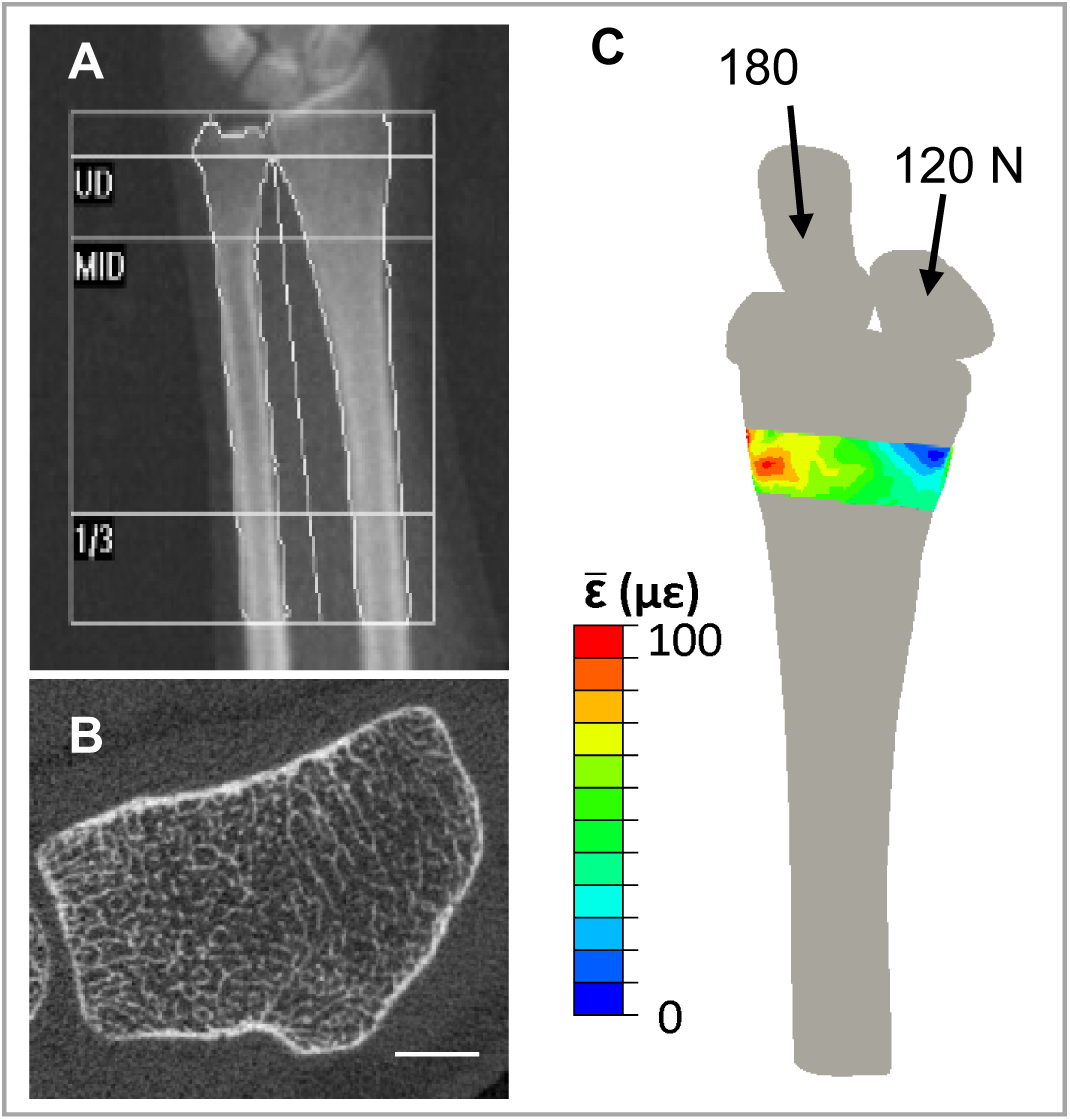
a) Representative forearm DXA scan including ultradistal (UD), Middle (MID) and 1/3 regions, and b) distal radius HRpQCT scan (scale bar 5 mm). c) Three-dimensional continuum FE model used to estimate energy equivalent strain 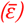 within the HRpQCT scanned region.

### 2.2 Anthropometrics and Loading Assessments

Height was measured using a wall-mounted stadiometer, and body mass was measured using an analog scale. Non-dominant grip strength was measured using a hydraulic hand-grip dynamometer (Baseline; White Plains, NJ) three times and averaged. Grip strength measurements were taken in a seated position with the elbow bent ninety degrees in flexion. Average daily calcium intake (mg/day) was estimated using a 10-item questionnaire that tallied weekly consumption of calcium-containing foods and beverages [40].

To estimate forearm loading due to physical activity, a site-specific arm bone loading index (*armBLI*) algorithm [41] was used to score activity histories. The *armBLI* algorithm scores activities based on the magnitude, rate, and frequency (days/week) of loads applied to the non-dominant arm as:

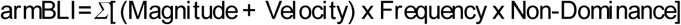

where the non-dominance multiplier corrects for activities loading the dominant arm preferentially. The multiplier is 0.33 for predominantly unilateral activities (e.g., tennis), 0.66 for somewhat unilateral activities (e.g. softball), and 1.0 for bilateral activities (e.g. gymnastics). For each individual, an overall score is calculated as the products of activity-specific training volumes and *armBLI* indices summed over all activities performed. For the present study, physical activity training volumes were generated using the validated Bone Loading History Questionnaire (BLHQ) [42], which was used to collect physical activity history in this group. Briefly, training volume is calculated as the product of years of participation, the seasons participated per year (fraction out of four), and a frequency score ranging from 1 to 4 reflecting training sessions per week (1=1-3 times per month, 2=1-2 times per week, 3=3-5 times per week, and 4=>5 times per week). To assess the relative importance of upper-extremity physical activity during different stages of development, separate mean annual scores (armBLI/year) were calculated for adolescent (age 10-18) and adult (age 19-current age) loading.

### 2.3 High-Resolution Peripheral Quantitative Computed Tomography

High-resolution peripheral quantitative computed tomography (HRpQCT; XtremeCT, Scanco Medical; Brüttisellen, Switzerland) scans of the distal radius in the non-dominant arm were performed according to the manufacturer’s standard *in vivo* scanning protocol (Figure 2b). The scans consisted of 110 slices with an isotropic voxel size of 82 µm, encompassing a 9.02 mm axial region beginning 9.5 mm proximal to a reference line placed at the distal endplate. All scans were performed by trained technicians, and daily and weekly quality control scans were performed. Each scan was graded for motion on a scale from 1 (no motion) to 5 (severe motion artifact) [43], and only scans scoring 3 or better were included in the analysis.

HRpQCT scans were analyzed using the manufacturer’s semi-automatic standard morphological [44] and cortical [45–48] analyses. Total vBMD (mgHA/cm^3^), trabecular vBMD (mgHA/cm^3^), total mean cross-sectional area (CSA; mm^2^), and trabecular number (mm^−1^) were calculated using the standard manufacturer’s analysis, and cortical vBMD (mgHA/cm^3^), cortical thickness (mm), and cortical porosity (%) were calculated using the dual-threshold method [45–48].

### 2.4 Continuum FE Modeling

Clinical resolution CT scans were used to construct three-dimensional continuum FE models including the distal articulating surface to simulate physiologic loading through the scaphoid and lunate. CT scans of the distal-most 12 cm of the non-dominant forearm were acquired using established methods with a transverse pixel size of 234 µm and slice thickness of 625 µm (BrightSpeed, GE Healthcare; Chicago, IL) [49]. A calibration phantom with known calcium hydroxyapatite equivalent concentrations was included for conversion from Hounsfield Units to apparent density.

Radius bone strain during the forearm loading task was estimated from FE models [17] simulating compressive loading of 300 N (approximately one half body-weight) through the palm of the hand (Figure 2c). Briefly, the radius, scaphoid and lunate were segmented from the CT scan using a 175 mgHA/cm^3^ density threshold in Mimics (Materialise; Leuven, Belgium) and converted to quadratic tetrahedral FE meshes with mean element edge length of 3 mm in 3Matic (Materialise; Leuven, Belgium). A 2 mm quadratic tetrahedral articular cartilage mesh was generated by dilating the radius in the transverse plane. Inhomogeneous, density-based material properties were assigned to the radius using an established density-elasticity relationship [50]. The scaphoid and lunate were modeled as incompressible, and cartilage as a neo-Hookean hyperelastic solid with *E*=10 MPa and *ν*=0.45. Loading was simulated in Abaqus 6.12 (Simulia; Providence, RI) by applying a ramped force through the centroids of the scaphoid (180 N) and lunate (120 N) towards the centroid of the fixed proximal radius. These methods have been validated using experimental testing of cadaveric specimens [17].

Energy equivalent strain 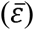 was calculated within the ultradistal region within the continuum FE model matching the HRpQCT scan region. Energy equivalent strain was selected as the primary FE outcome because it has been previously related to radius bone adaptation [49]. This scalar quantity represents the total work done on the bone tissue, provided by the multi-axial stress-strain state:

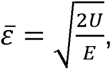

where E is the elastic modulus, and U is the strain energy density calculated as:

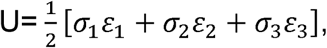

where σ_n_ and ε_n_ are the principal stress and strain components, respectively. Mean energy equivalent strain within the region corresponding to the HRpQCT-scanned region was identified using a custom Matlab script that implemented a mutual information image registration algorithm considering pixel intensities. A laboratory precision study yielded mean rotation errors of 0.47±0.38°, 0.46±0.41°, and 0.32±0.24° in the x, y, and z directions, respectively, for a similar data set [51].

### 2.5 Statistical Analysis

The normality of each measured variable was assessed by visual inspection of histogram distributions. To assess the ability of DXA-based measures to predict FE strain, a power regression model was constructed with ultradistal aBMD as the independent variable and mean energy equivalent strain as the dependent variable. Power regression was selected based on previous studies characterizing the relationship between bone density and mechanical properties [52]. Correlation and multiple regression analyses were used to identify intrinsic and modifiable factors that affect bone strain and distal radius microstructure. Pearson and Spearman correlation coefficients were calculated between subject characteristics, FE-strain, and HRpQCT parameters with normal and non-normal distributions, respectively. A series of hierarchical linear regression models were fitted for each structure and strain variable, with age and height entered as intrinsic covariates and body mass, grip strength, and loading scores included as extrinsic, modifiable predictors. Covariates were added as a first block of independent variables, and then a single modifiable factor was entered in a second block, allowing the total variance explained by the intrinsic factors as a group and the predictive capability of each loading factor to be determined. The overall model residuals were visually inspected for normality and homoscedasticity using a plot of residuals versus predicted values. An alpha level of 0.05 was used to detect significance. All statistical analyses were performed using SPSS v22.0.

## 3. Results

### 3.1 Subject Characteristics

Descriptive statistics, presented as means and standard deviations, are summarized in Table 1. Ten enrolled subjects were excluded from analyses due to incomplete physical activity data (n=3) or HRpQCT motion artifact (n=7). Thus, all results are reported for the seventy-two subjects for whom complete data were available. Daily calcium intake was below the average intake reported for women ages 19-50 in the United States [53], while grip strength was similar to previously reported values for young adult women [54,55]. Correlation coefficients between predictors and bone structure and strain parameters are provided in Table 2. Mean energy equivalent strain within the distal region was significantly correlated several HRpQCT parameters, DXA aBMD, and body mass.

**Table 1:** Descriptive statistics for all subjects (n=72)

| Subject Characteristics | Mean | SD |
| --- | --- | --- |
| Age (years) | 28.3 | 5.3 |
| Body Mass (kg) | 63.6 | 8.6 |
| Height (cm) | 164.6 | 6.9 |
| ND Grip Strength (kg) | 26.4 | 5.2 |
| Vit D (ng/mL) | 32 | 9 |
| Daily calcium intake (mg/day) | 682 | 400 |
| Adolescent Loading Score (armBLI/year) | 50 | 48 |
| Adult Loading Score (armBLI/year) | 54 | 47 |

*Vit D* Serum Vitamin D level, *ArmBLI* Arm Bone Loading Index

**Table 2:**
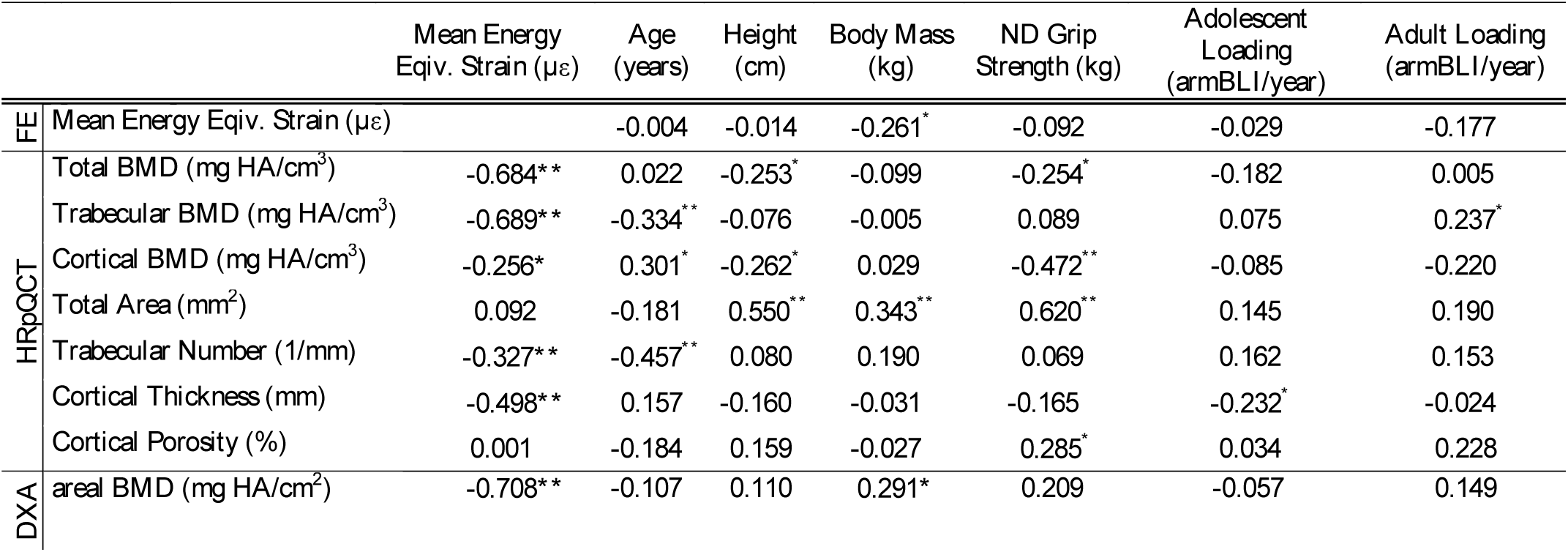
Correlation coefficients between subject characteristics, bone structure, and strain parameters. *p<0.05, **p<0.01

### 3.2 Prediction of FE-Estimated Bone Strain and HRpQCT Microstructure

Results of the nonlinear power regression between DXA aBMD and mean energy equivalent strain within the ultradistal region are presented in Figure 3. Areal BMD explained 51.47% of the variability in strain, with higher density values associated with lower strains under a given load.

**Figure 3:**
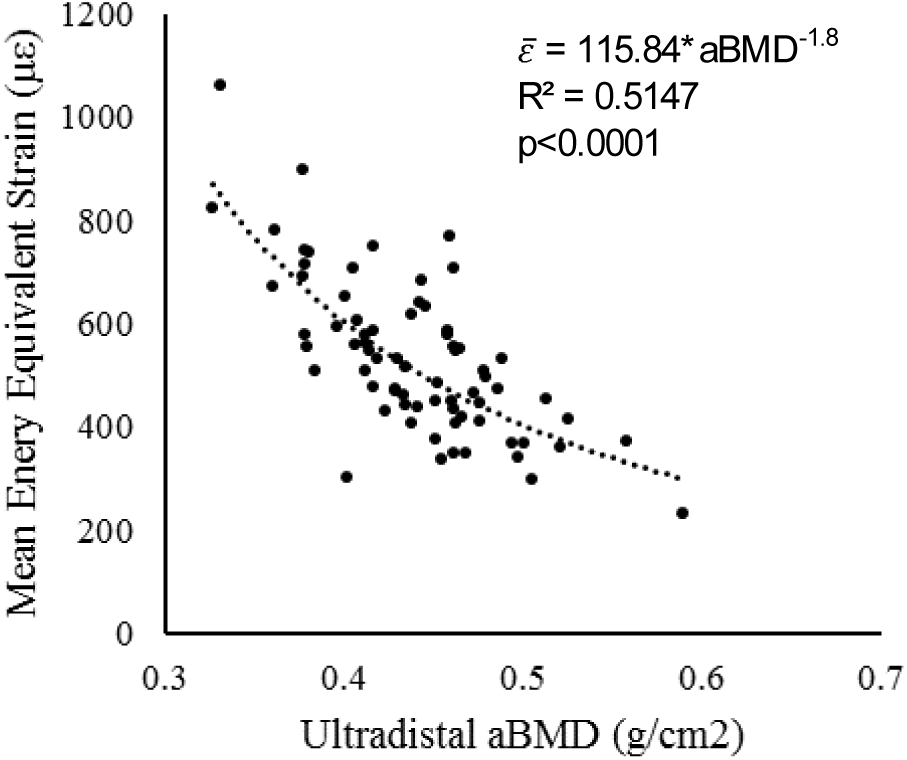
Mean energy equivalent strain within the ultradistal region matching the volume scanned with HRpQCT versus areal bone mineral density measured using DXA within the standard ultradistal site.

Mean and standard deviations for all bone parameters, as well as the corresponding hierarchical regression results, are presented in Tables 3 and 4. Mean values for HRpQCT-measured parameters agree well with those reported for young adult women [56]. Energy equivalent strain was not significantly predicted by age or height. Adding body mass to the model significantly improved the prediction of strain, explaining an additional 10.0% of the variance (p=0.008).

**Table 3:**
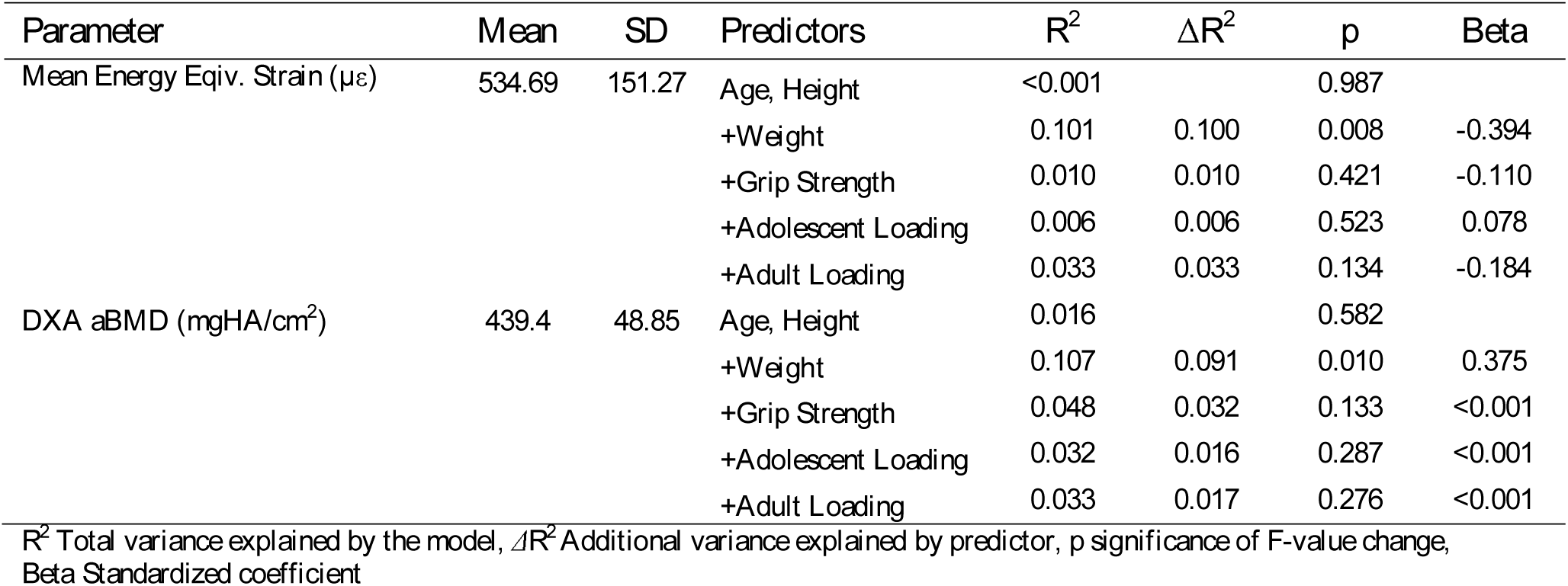
HRpQCT parameter values (mean±SD) and hierarchical linear regression results

| Parameter | Mean | SD | Predictors | $R^2$ | $\Delta R^2$ | p | Beta |
| --- | --- | --- | --- | --- | --- | --- | --- |
| Mean Energy Eqiv. Strain ($\mu\epsilon$) | 534.69 | 151.27 | Age, Height | <0.001 | | 0.987 | |
|  |  |  | **+Weight** | **0.101** | **0.100** | **0.008** | **-0.394** |
|  |  |  | +Grip Strength | 0.010 | 0.010 | 0.421 | -0.110 |
|  |  |  | +Adolescent Loading | 0.006 | 0.006 | 0.523 | 0.078 |
|  |  |  | +Adult Loading | 0.033 | 0.033 | 0.134 | -0.184 |
| DXA aBMD (mgHA/cm<sup>2</sup>) | 439.4 | 48.85 | Age, Height | 0.016 |  | 0.582 |  |
|  |  |  | **+Weight** | **0.107** | **0.091** | **0.010** | **0.375** |
|  |  |  | +Grip Strength | 0.048 | 0.032 | 0.133 | <0.001 |
|  |  |  | +Adolescent Loading | 0.032 | 0.016 | 0.287 | <0.001 |
|  |  |  | +Adult Loading | 0.033 | 0.017 | 0.276 | <0.001 |

$R^2$ Total variance explained by the model, $\Delta R^2$ Additional variance explained by predictor, $p$ significance of F-value change, $Beta$ Standardized coefficient

**Table 4:** HRpQCT parameter values (mean±SD) and hierarchical linear regression results

| Parameter | Mean | SD | Predictors | R<sup>2</sup> | ΔR<sup>2</sup> | p | Beta |
| --- | --- | --- | --- | --- | --- | --- | --- |
| Total BMD (mgHA/cm<sup>3</sup>) | 298.23 | 51.53 | Age, Height | 0.064 |  | 0.101 |  |
|  |  |  | +Body Mass | 0.067 | 0.003 | 0.640 | 0.068 |
|  |  |  | +Grip Strength | 0.088 | 0.024 | 0.184 | -0.175 |
|  |  |  | +Adolescent Loading | 0.068 | 0.004 | 0.584 | -0.065 |
|  |  |  | +Adult Loading | 0.076 | 0.012 | 0.360 | 0.110 |
| Trabecular BMD (mgHA/cm<sup>3</sup>) | 162.86 | 30.03 | **Age, Height** | **0.096** |  | **0.031** |  |
|  |  |  | + Body Mass | 0.110 | 0.014 | 0.309 | 0.146 |
|  |  |  | +Grip Strength | 0.122 | 0.026 | 0.164 | 0.181 |
|  |  |  | +Adolescent Loading | 0.101 | 0.005 | 0.525 | 0.074 |
|  |  |  | **+Adult Loading** | **0.167** | **0.071** | **0.019** | **0.272** |
| Cortical BMD (mgHA/cm<sup>3</sup>) | 969.24 | 44.06 | **Age, Height** | **0.119** |  | **0.013** |  |
|  |  |  | + Body Mass | 0.147 | 0.028 | 0.142 | 0.207 |
|  |  |  | **+Grip Strength** | **0.289** | **0.170** | **<0.001** | **-0.464** |
|  |  |  | +Adolescent Loading | 0.120 | 0.001 | 0.749 | -0.037 |
|  |  |  | +Adult Loading | 0.139 | 0.020 | 0.215 | -0.144 |
| Total Area (mm<sup>2</sup>) | 274.23 | 49.23 | **Age, Height** | **0.316** |  | **<0.001** |  |
|  |  |  | + Body Mass | 0.321 | 0.005 | 0.493 | 0.086 |
|  |  |  | **+Grip Strength** | **0.494** | **0.179** | **<0.001** | **0.477** |
|  |  |  | +Adolescent Loading | 0.316 | <0.001 | 0.945 | -0.007 |
|  |  |  | +Adult Loading | 0.325 | 0.009 | 0.354 | 0.095 |
| Trabecular Number (1/mm) | 2.00 | 0.26 | **Age, Height** | **0.174** |  | **0.001** |  |
|  |  |  | **+ Body Mass** | **0.249** | **0.076** | **0.011** | **0.342** |
|  |  |  | +Grip Strength | 0.178 | 0.004 | 0.564 | 0.072 |
|  |  |  | +Adolescent Loading | 0.195 | 0.022 | 0.179 | 0.148 |
|  |  |  | +Adult Loading | 0.198 | 0.024 | 0.158 | 0.158 |
| Cortical Thickness (mm) | 0.77 | 0.15 | Age, Height | 0.047 |  | 0.189 |  |
|  |  |  | + Body Mass | 0.049 | 0.002 | 0.738 | 0.049 |
|  |  |  | +Grip Strength | 0.060 | 0.013 | 0.337 | -0.128 |
|  |  |  | +Adolescent Loading | 0.072 | 0.025 | 0.178 | -0.159 |
|  |  |  | +Adult Loading | 0.048 | 0.001 | 0.846 | 0.024 |
| Cortical Porosity (%) | 1.20 | 0.67 | Age, Height | 0.050 |  | 0.169 |  |
|  |  |  | + Body Mass | 0.091 | 0.041 | 0.084 | -0.252 |
|  |  |  | **+Grip Strength** | **0.106** | **0.056** | **0.043** | **0.267** |
|  |  |  | +Adolescent Loading | 0.050 | <0.001 | 0.946 | 0.008 |
|  |  |  | **+Adult Loading** | **0.133** | **0.083** | **0.013** | **0.294** |

*R*<sup>2</sup> Total variance explained by the model, Δ*R*<sup>2</sup> Additional variance explained by predictor, *p* significance of F-value change, *Beta* Standardized coefficient

Looking at HRpQCT parameters, age and height accounted for 9.6% of the variance in trabecular BMD (p=0.031) as intrinsic factors, and adding adult loading score to the model accounted for an additional 7.1% of the variance (p=0.019). Intrinsic factors alone explained 11.9% of the variance in cortical BMD (p=0.013), and adding grip strength to the model explained an additional 17.0% of the variance (p<0.001). Total cross sectional area was strongly predicted by age and height, which explained 31.6% of the variance (p<0.001). Adding grip strength to the model significantly improved the prediction of total area, explaining an additional 17.9% of the variance (p<0.001). Intrinsic factors alone explained 17.4% of the variance in trabecular number (p=0.001), and body mass accounted for an additional 7.6% percent of the variance (p=0.011). Cortical porosity was not significantly predicted by intrinsic factors alone, but adding either grip strength or adult loading score improved model predictions by 5.6% (p=0.043) and 8.3% (p=0.013), respectively. None of the models predicting total BMD or cortical thickness were significant.

## 4. Discussion

Our primary purpose was to quantify the inter-individual variability in radius microstructure and FE-estimated strain explained by site-specific mechanical loading history as described by grip strength, physical activity history, and body weight. Higher grip strength predicted greater cross-sectional area and lower cortical density related to greater cortical porosity. Similar trends were seen in individuals with higher levels of site-specific adult loading, who tended to have more porous cortices and greater trabecular density. Neither grip strength nor adult loading significantly predicted strain, suggesting that differences in cortical density and cross-sectional area may compensate for each other with respect to whole-radius mechanics. Finally, greater body mass predicted higher trabecular number and lower ultradistal strain. This suggests that within the normal BMI range, greater body mass is associated with improved mechanical behavior (i.e. lower strains under a given load), which may be attributed to more interconnected trabeculae supporting the distal region. As covariate factors, age and height were significant predictors of trabecular number, trabecular and cortical vBMD, and cross-sectional area but not total vBMD, cortical thickness, cortical porosity or bone strain. Taken together, these results suggest that meaningful differences in bone morphology and mechanical behavior can be predicted by measures of site-specific mechanical loading.

Secondly, using non-linear regression, we determined whether variability in FE-estimated radius strain was captured by aBMD, a clinically relevant measure of bone density and fracture risk. While aBMD and FE strain are associated, over forty-eight percent of the variability in strain was left unexplained by its relationship with density. Thus, while aBMD may help identify those most in need of bone-building interventions, it does not fully describe the mechanical behavior of bone tissue under loading, which is critical to predicting cell-driven adaptation.

In the current study, the contribution of upper extremity mechanical loading was considered through the inclusion of body mass, grip strength, and questionnaire-based physical activity scores. These measures are to some extent related, as more active individuals may have greater muscle mass, which affects both grip strength and body mass. However, each has been previously related to bone quality and may characterize different aspects or modes of loading. For example, individuals with greater body mass experience larger compressive loads during weight-bearing exercises, to which bone adapts. This is consistent with the observation that heavier individuals experienced lower-magnitude strains for a given compressive force, indicating stronger bone. This also supports the previously reported effects of body mass on lower-extremity mechanical loading [57], aBMD [58], and fracture risk [27]. Grip strength is a functionally useful measure of muscle mass and strength, and has been associated with bone density, macrostructure, and strength using peripheral QCT [24–26]. The relationship between muscle mass and bone quality is complex. Individuals may be genetically predisposed to having larger muscles and bones, or may build muscle over time, which drives bone adaptation to increasing forces. In the current study, grip strength was positively related to cross sectional area but not strain, suggesting that grip strength may describe global body size rather than adaptation to specific loads.

Physical activity during growth and adulthood has been associated with improvements to bone structure [59]. However, there is a lack of consensus whether loading during adolescence or early adulthood are more significant in determining peak bone mass [60–62]. We found that adolescent loading did not significantly contribute to the prediction of any bone structural or parameter, while adult loading was associated with favorable trabecular vBMD. Variations between previous and the current results may be related to differences in questionnaires or anatomic sites. As opposed to other skeletal loading questionnaires, the *armBLI* scores activities based on the magnitude and frequency of forearm loading rather than using ground reaction forces [63] or estimations of loading at the hip and spine [42]. The relationship between loading and structure may also be site-specific, especially considering the differences in habitual loading between the upper and lower extremities.

Cortical porosity was significantly predicted by grip strength and adult loading score. However, correlation coefficients in both cases were negative, indicating that more active individuals with greater muscle mass have more porous cortices. This is somewhat surprising, as increased cortical porosity is associated with diminished structural integrity and increased fracture risk in older individuals [48]. However, increased cortical porosity in this younger population may reflect more active remodeling units rather than degradation, driven by adaptation to increased applied loading. This finding is consistent with a previous study in females ages 14-21 that found significantly higher cortical porosity in both amenorrheic and eumenorrheic athletes versus non-athletes, despite higher HR-pQCT FE-derived stiffness in eumenorrheic athletes [64]. The relationship between grip strength and porosity, however, should be interpreted in the context of the HR-pQCT scanning protocol that we used, which defined the scan region a fixed, not relative, 9.5 mm distance from the distal endplate. As a result, the scanned region was shifted distally in individuals with longer forearms, potentially biasing results towards increased cross sectional area and cortical porosity [65]. Recent work has highlighted the potential advantage of defining scan regions relative to limb length, which may reduce the variability in results associated with body size [65]. As height and grip strength are positively related [66], it is possible that the positive relationships between grip strength, total CSA, and cortical porosity observed here may be, in part, due to individuals with higher grip strength having longer forearms and thus more distal scan regions. However, in the current analysis, all regression models were statistically controlled for height to mitigate this effect. Our results are further strengthened by the lack of significant relationship between cortical porosity, age, and height.

The current study is not without limitations. Our sample size was relatively small, and subjects were recruited as part of a longitudinal study with inclusion criteria developed for the evaluation of a loading intervention. To target individuals who would most likely benefit from new loading, anyone already regularly participating in activities involving frequent, high impact loading of the upper extremities was excluded. Additionally, only women with a DXA total forearm T-score falling within the range −2.5 to 1.0 were included. Therefore, the current results cannot be generalized to women with extreme levels of upper-extremity loading, those with bone mass below the expected range for their age, or those with T-score more than 1.0 SD above the population mean. Additionally, there may have been limitations in applying the *armBLI* algorithm to adult women with retrospective rather than prospective, calendar-based training histories. The accuracy with which adolescent activity was recalled may have been limited and introduced additional variability, contributing to the lack of significant predictions by adolescent loading. Further, the *armBLI* was validated against DXA areal density measurements [41] rather than volumetric structure or FE-derived strain. Considering these differences, a more rigorous validation of the *armBLI* may be required in adult women using CT-based measurements.

In summary, we have shown that individuals with higher levels of adult physical activity, grip strength, and body mass tend to have favorable bone microstructure structure. Women with higher body mass within a normal BMI range also had lower levels of strain under a given force, suggestive of adaptation to increased loads during functional activities. Additionally, we have explored the relationships between clinical measures of bone quality, showing that the current gold-standard, DXA aBMD, does not capture the wide range of strains experienced during typical physiologic loading. Overall, these results suggest the importance of engaging in bone-building behaviors in adulthood, and contribute to the systematic design of prescribed loading interventions to better address the growing incidence of osteoporotic fracture.

## Acknowledgements

This work was supported by the National Institutes of Health [grant number R01AR063691 (KLT)]; and the National Science Foundation [grant numbers DGE1106756 (MM), DGE1144804 (MM)]. The content is solely the responsibility of the authors and does not necessarily represent the official views of the funding agencies.

## References

[1] O.O. Babatunde, J.J. Forsyth, C.J. Gidlow, A meta-analysis of brief high-impact exercises for enhancing bone health in premenopausal women, Osteoporos. Int. 23 (2012) 109–119. doi:10.1007/s00198-011-1801-0.

[2] G.A. Kelley, K.S. Kelley, W.M. Kohrt, Exercise and bone mineral density in premenopausal women: A meta-analysis of randomized controlled trials, Int. J. Endocrinol. 2013 (2013). doi:10.1155/2013/741639.

[3] M.M.S. James, S. Carroll, Effects of different impact exercise modalities on bone mineral density in premenopausal women: A meta-analysis, J. Bone Miner. Metab. 28 (2010) 251–267. doi:10.1007/s00774-009-0139-6.

[4] M. Martyn-St James, S. Carroll, Progressive High-Intensity Resistance Training and Bone Mineral Density Changes Among Premenopausal Women, Sport. Med. 36 (2006) 683–704. doi:10.2165/00007256-200636080-00005.

[5] F. Cosman, S.J. de Beur, M.S. LeBoff, E.M. Lewiecki, B. Tanner, S. Randall, R. Lindsay, Clinician’s Guide to Prevention and Treatment of Osteoporosis, Osteoporos. Int. 25 (2014) 2359–2381. doi:10.1007/s00198-014-2794-2.

[6] K.F. Janz, D.Q. Thomas, M.A. Ford, S.M. Williams, Top 10 Research Questions Related to Physical Activity and Bone Health in Children and Adolescents, Res. Q. Exerc. Sport. 86 (2015) 5–12. doi:10.1080/02701367.2015.991265.

[7] S.J. Warden, R.K. Fuchs, C.H. Turner, Steps for targeting exercise towards the skeleton to increase bone strength., Eura. Medicophys. 40 (2004) 223–32.

[8] C.T. Rubin, L.E. Lanyon, Regulation of bone mass by mechanical strain magnitude, Calcif. Tissue Int. 37 (1985) 411–417.

[9] Y.F. Hsieh, C.H. Turner, Effects of loading frequency on mechanically induced bone formation., J. Bone Miner. Res. 16 (2001) 918–924. doi:10.1359/jbmr.2001.16.5.918.

[10] C.T. Rubin, L.E. Lanyon, Regulation of Bone Formation by Applied Dynamic Loads, J. Bone Jt. Surg. 66A (1984) 397–402.

[11] Y. Umemura, T. Ishiko, T. Yamauchi, M. Kurono, S. Mashiko, Five jumps per day increase bone mass and breaking force in rats, J. Bone Miner. Res. 12 (1997) 1480–1485. doi:10.1359/jbmr.1997.12.9.1480.

[12] D. Webster, F.A. Schulte, F.M. Lambers, G. Kuhn, R. Müller, Strain energy density gradients in bone marrow predict osteoblast and osteoclast activity: A finite element study, J. Biomech. 48 (2015) 866–874. doi:10.1016/j.jbiomech.2014.12.009.

[13] F.M. Lambers, G. Kuhn, C. Weigt, K.M. Koch, F.A. Schulte, R. Müller, Bone adaptation to cyclic loading in murine caudal vertebrae is maintained with age and directly correlated to the local micromechanical environment, J. Biomech. 48 (2015) 1179–1187. doi:10.1016/j.jbiomech.2014.11.020.

[14] Z. Földhazy, A. Arndt, C. Milgrom, A. Finestone, I. Ekenman, Exercise-induced strain and strain rate in the distal radius, J Bone Jt. Surg Br. 87–b (2005) 261–266. doi:10.1302/0301-620X.87B2.14857.

[15] L.E. Lanyon, W.G.J. Hampson, A.E. Goodship, J.S. Shah, Bone Deformation Recorded in vivo from Strain Gauges Attached to the Human Tibial Shaft, Acta Orthop. Scand. 46 (1975) 256–268. doi:10.3109/17453677508989216.

[16] K.L. Troy, W.B. Edwards, V.A. Bhatia, M. Lou Bareither, In vivo loading model to examine bone adaptation in humans: A pilot study, J. Orthop. Res. 31 (2013) 1406–1413. doi:10.1002/jor.22388.

[17] V.A. Bhatia, W.B. Edwards, K.L. Troy, Predicting surface strains at the human distal radius during an in vivo loading task – Finite element model validation and application, J. Biomech. 47 (2014) 2759–2765. doi:10.1016/j.jbiomech.2014.04.050.

[18] C.M. Court-Brown, B. Caesar, Epidemiology of adult fractures: A review, Injury. 37 (2006) 691–697. doi:10.1016/j.injury.2006.04.130.

[19] A. Laib, H.J. Hauselmann, P. Ruegsegger, In vivo high resolution 3D-QCT of the human forearm, Technol. Heal. Care. 6 (1998) 329–337.

[20] M. Pasanen, S. Kontulainen, P. Kannus, H. Haapasalo, H. Sieva, Good Maintenance of Exercise-Induced Bone Gain with Decreased Training of Female Tennis and Squash Players□: A Prospective 5-Year Follow-Up Study of Young and Old Starters and Controls, 16 (2001) 195–201.

[21] R. Nikander, P. Kannus, T. Rantalainen, K. Uusi-Rasi, A. Heinonen, H. Sievänen, Cross-sectional geometry of weight-bearing tibia in female athletes subjected to different exercise loadings, Osteoporos. Int. 21 (2010) 1687–1694. doi:10.1007/s00198-009-1101-0.

[22] D. Courteix, E. Lespessailles, S.L. Peres, P. Obert, P. Germain, C.L. Benhamou, Effect of physical training on bone mineral density in prepubertal girls: a comparative study between impact-loading and non-impact-loading sports, Osteoporos Int. 8 (1998) 152–158. doi:10.1007/bf02672512.

[23] T.A. Scerpella, B. Bernardoni, S. Wang, P.J. Rathouz, Q. Li, J.N. Dowthwaite, Site-specific, adult bone benefits attributed to loading during youth: A preliminary longitudinal analysis, Bone. 85 (2016) 148–159. doi:10.1016/j.bone.2016.01.020.

[24] Y. Hasegawa, P. Schneider, C. Reiners, Age, sex, and grip strength determine architectural bone parameters assessed by peripheral quantitative computed tomography (pQCT) at the human radius, J. Biomech. 34 (2001) 497–503. doi:10.1016/S0021-9290(00)00211-6.

[25] A.L. Lorbergs, J.P. Farthing, A.D.G. Baxter-Jones, S.A. Kontulainen, Forearm muscle size, strength, force, and power in relation to pQCT-derived bone strength at the radius in adults, Appl. Physiol. Nutr. Metab. 36 (2011) 618–625. doi:10.1139/h11-065.

[26] K.G. Greenway, J.W. Walkley, P.A. Rich, Relationships between self-reported lifetime physical activity, estimates of current physical fitness, and aBMD in adult premenopausal women, Arch. Osteoporos. 10 (2015). doi:10.1007/s11657-015-0239-y.

[27] S. Morin, J.F. Tsang, W.D. Leslie, Weight and body mass index predict bone mineral density and fractures in women aged 40 to 59 years, Osteoporos. Int. 20 (2009) 363–370. doi:10.1007/s00198-008-0688-x.

[28] C.G. Gjesdal, J.I. Halse, G.E. Eide, J.G. Brun, G.S. Tell, Impact of lean mass and fat mass on bone mineral density: The Hordaland Health Study, Maturitas. 59 (2008) 191–200. doi:10.1016/j.maturitas.2007.11.002.

[29] L.A. Rubin, G.A. Hawker, V.D. Peltekova, L.J. Fielding, R. Ridout, D.E. Cole, Determinants of peak bone mass: clinical and genetic analyses in a young female Canadian cohort., J. Bone Miner. Res. 14 (1999) 633–43. doi:10.1359/jbmr.1999.14.4.633.

[30] A.Y.Y. Ho, A.W.C. Kung, Determinants of peak bone mineral density and bone area in young women, J. Bone Miner. Metab. 23 (2005) 470–475. doi:10.1007/s00774-005-0630-7.

[31] D.T. Felson, Y. Zhang, M.T. Hannan, J.J. Anderson, Effects of weight and body mass index on bone mineral density in men and women: The framingham study, J. Bone Miner. Res. 8 (1993) 567–573. doi:10.1002/jbmr.5650080507.

[32] E. Lespessailles, R. Hambli, S. Ferrari, Osteoporosis drug effects on cortical and trabecular bone microstructure: a review of HR-pQCT analyses, Bonekey Rep. 5 (2016) 1–8. doi:10.1038/bonekey.2016.59.

[33] R. Hambli, Micro-CT finite element model and experimental validation of trabecular bone damage and fracture, Bone. 56 (2013) 363–374. doi:10.1016/j.bone.2013.06.028.

[34] E.M. Stein, X.S. Liu, T.L. Nickolas, A. Cohen, V. Thomas, D.J. McMahon, C. Zhang, P.T. Yin, F. Cosman, J. Nieves, X.E. Guo, E. Shane, Abnormal microarchitecture and reduced stiffness at the radius and tibia in postmenopausal women with fractures, J. Bone Miner. Res. 25 (2010) 2296–2305. doi:10.1002/jbmr.152.

[35] M.H. Edwards, D.E. Robinson, K.A. Ward, M.K. Javaid, K. Walker-Bone, C. Cooper, E.M. Dennison, Cluster analysis of bone microarchitecture from high resolution peripheral quantitative computed tomography demonstrates two separate phenotypes associated with high fracture risk in men and women, Bone. 88 (2016) 131–137. doi:10.1016/j.bone.2016.04.025.

[36] E. Sornay-Rendu, S. Boutroy, F. Duboeuf, R.D. Chapurlat, Bone Microarchitecture Assessed by HR-pQCT as Predictor of Fracture Risk in Postmenopausal Women: The OFELY Study, J. Bone Miner. Res. 32 (2017) 1243–1251. doi:10.1002/jbmr.3105.

[37] M. Nilsson, C. Ohlsson, D. Sundh, D. Mellström, M. Lorentzon, Association of physical activity with trabecular microstructure and cortical bone at distal tibia and radius in young adult men, J. Clin. Endocrinol. Metab. 95 (2010) 2917–2926. doi:10.1210/jc.2009-2258.

[38] M. Nilsson, C. Ohlsson, D. Mellström, M. Lorentzon, Sport-specific association between exercise loading and the density, geometry, and microstructure of weight-bearing bone in young adult men, Osteoporos. Int. 24 (2013) 1613–1622. doi:10.1007/s00198-012-2142-3.

[39] J.D. Schipilow, H.M. Macdonald, A.M. Liphardt, M. Kan, S.K. Boyd, Bone micro-architecture, estimated bone strength, and the muscle-bone interaction in elite athletes: An HR-pQCT study, Bone. 56 (2013) 281–289. doi:10.1016/j.bone.2013.06.014.

[40] P.C. Questionnaire, Patient Calcium Questionnaire GETTING ENOUGH CALCIUM□? Information about Calcium, (n.d.).

[41] J.N. Dowthwaite, K.A. Dunsmore, N.M. Gero, A.O. Burzynski, C.A. Sames, P.F. Rosenbaum, T.A. Scerpella, Arm bone loading index predicts DXA musculoskeletal outcomes in two samples of post-menarcheal girls, J. Musculoskelet. Neuronal Interact. 15 (2015) 358–371.

[42] S.H. Dolan, D.P. Williams, B.E. Ainsworth, J.M. Shaw, Development and reproducibility of the bone loading history questionnaire, Med. Sci. Sports Exerc. 38 (2006) 1121–1131. doi:10.1249/01.mss.0000222841.96885.a8.

[43] J.B. Pialat, A.J. Burghardt, M. Sode, T.M. Link, S. Majumdar, Visual grading of motion induced image degradation in high resolution peripheral computed tomography: Impact of image quality on measures of bone density and micro-architecture, Bone. 50 (2012) 111–118. doi:10.1016/j.bone.2011.10.003.

[44] J.A. MacNeil, S.K. Boyd, Accuracy of high-resolution peripheral quantitative computed tomography for measurement of bone quality., Med. Eng. Phys. 29 (2007) 1096–1105. doi:10.1016/j.medengphy.2006.11.002.

[45] H.R. Buie, G.M. Campbell, R.J. Klinck, J.A. MacNeil, S.K. Boyd, Automatic segmentation of cortical and trabecular compartments based on a dual threshold technique for in vivo micro-CT bone analysis, Bone. 41 (2007) 505–515. doi:10.1016/j.bone.2007.07.007.

[46] A.J. Burghardt, H.R. Buie, A. Laib, S. Majumdar, S.K. Boyd, Reproducibility of direct quantitative measures of cortical bone microarchitecture of the distal radius and tibia by HR-pQCT, Bone. 47 (2010) 519–528. doi:10.1016/j.bone.2010.05.034.

[47] A.J. Burghardt, G.J. Kazakia, S. Ramachandran, T.M. Link, S. Majumdar, Age and Gender Related Differences in the Geometric Properties and Biomechanical Significance of Intra-Cortical Porosity in the Distal Radius and Tibia, J. Bone Miner. Res. 25 (2009) 983–993. doi:10.1359/jbmr.091104.

[48] K.K. Nishiyama, H.M. Macdonald, H.R. Buie, D.A. Hanley, S.K. Boyd, Postmenopausal Women With Osteopenia Have Higher Cortical Porosity and Thinner Cortices at the Distal Radius and Tibia Than Women With Normal aBMD: An In Vivo HR-pQCT Study, J. Bone Miner. Res. 25 (2009) 882–890. doi:10.1359/jbmr.091020.

[49] V.A. Bhatia, W. Brent Edwards, J.E. Johnson, K.L. Troy, Short-Term Bone Formation is Greatest Within High Strain Regions of the Human Distal Radius: A Prospective Pilot Study, J. Biomech. Eng. 137 (2015) 11001–1–5. doi:10.1115/1.4028847.

[50] E.F. Morgan, H.H. Bayraktar, T.M. Keaveny, Trabecular bone modulus-density relationships depend on anatomic site, J. Biomech. 36 (2003) 897–904. doi:10.1016/S0021-9290(03)00071-X.

[51] J.E. Johnson, K.L. Troy, Validation of a new multiscale finite element analysis approach at the distal radius, Med. Eng. Phys. 44 (2017) 16–24. doi:10.1016/j.medengphy.2017.03.005.

[52] B. Helgason, E. Perilli, E. Schileo, F. Taddei, S. Brynjólfsson, M. Viceconti, Mathematical relationships between bone density and mechanical properties: A literature review, Clin. Biomech. 23 (2008) 135–146. doi:10.1016/j.clinbiomech.2007.08.024.

[53] R.L. Bailey, K.W. Dodd, J. a Goldman, J.J. Gahche, J.T. Dwyer, A.J. Moshfegh, C.T. Sempos, M.F. Picciano, Estimation of Total Usual Calcium and Vitamin D Intakes in the United States 1 – 3, J. Nutr. (2010) 817–822. doi:10.3945/jn.109.118539.817.

[54] S.L. Wong, Grip strength reference values for Canadians aged 6 to 79: Canadian Health Measures Survey, 2007 to 2013, (2016).

[55] N.M. Massy-Westropp, T.K. Gill, A.W. Taylor, R.W. Bohannon, C.L. Hill, Hand Grip Strength: age and gender stratified normative data in a population-based study., BMC Res. Notes. 4 (2011) 127. doi:10.1186/1756-0500-4-127.

[56] L.A. Burt, H.M. Macdonald, D.A. Hanley, S.K. Boyd, Bone microarchitecture and strength of the radius and tibia in a reference population of young adults: an HR-pQCT study., Arch. Osteoporos. 9 (2014) 183. doi:10.1007/s11657-014-0183-2.

[57] M. Lou Bareither, K.L. Troy, M.D. Grabiner, Bone mineral density of the proximal femur is not related to dynamic joint loading during locomotion in young women, Bone. 38 (2006) 125–129. doi:10.1016/j.bone.2005.07.003.

[58] S. Gnudi, E. Sitta, N. Fiumi, Relationship between body composition and bone mineral density in women with and without osteoporosis: Relative contribution of lean and fat mass, J. Bone Miner. Metab. 25 (2007) 326–332. doi:10.1007/s00774-007-0758-8.

[59] T. Kato, T. Yamashita, S. Mizutani, A. Honda, M. Matumoto, Y. Umemura, Adolescent exercise associated with long-term superior measures of bone geometry: a cross-sectional DXA and MRI study., Br. J. Sports Med. 43 (2009) 932–935. doi:10.1136/bjsm.2008.052308.

[60] K.J. MacKelvie, K.M. Khan, H. a McKay, Is there a critical period for bone response to weight-bearing exercise in children and adolescents? a systematic review., Br. J. Sports Med. 36 (2002) 250–257; discussion 257. doi:10.1136/bjsm.36.4.250.

[61] R.M. Daly, S.L. Bass, Lifetime sport and leisure activity participation is associated with greater bone size, quality and strength in older men, Osteoporos. Int. 17 (2006) 1258–1267. doi:10.1007/s00198-006-0114-1.

[62] S. Bass, G. Pearce, M. Bradney, E. Hendrich, P.D. Delmas, A. Harding, E. Seeman, Exercise Before Puberty May Confer Residual Benefits in Bone Density in Adulthood: Studies in Active Prepubertal and Retired Female Gymnasts, J. Bone Miner. Res. 13 (1998) 500–507. doi:10.1359/jbmr.1998.13.3.500.

[63] B.K. Weeks, B.R. Beck, The BPAQ: A bone-specific physical activity assessment instrument, Osteoporos. Int. 19 (2008) 1567–1577. doi:10.1007/s00198-008-0606-2.

[64] K.E. Ackerman, M. Putman, G. Guereca, A.P. Taylor, L. Pierce, D.B. Herzog, A. Klibanski, M. Bouxsein, M. Misra, Cortical microstructure and estimated bone strength in young amenorrheic athletes, eumenorrheic athletes and non-athletes, Bone. 51 (2012) 680–687. doi:10.1016/j.bone.2012.07.019.

[65] A. Ghasem-Zadeh, A. Burghardt, X.F. Wang, S. Iuliano, S. Bonaretti, M. Bui, R. Zebaze, E. Seeman, Quantifying sex, race, and age specific differences in bone microstructure requires measurement of anatomically equivalent regions, Bone. 101 (2017) 206–213. doi:10.1016/j.bone.2017.05.010.

[66] J.A. Balogun, A.A. Akinloye, S.A. Adenlola, Grip strength as a function of age, height, body weight and quetelet index, Physiother. Theory Pract. 7 (1991) 111–119. doi:10.3109/09593989109106961.

